# *Rhizobium rhizogenes* A4-derived strains mediate hyper-efficient transient gene expression in *Nicotiana benthamiana* and other solanaceous plants

**DOI:** 10.1101/2024.11.30.626145

**Authors:** Juan Carlos Lopez-Agudelo, Foong-Jing Goh, Sopio Tchabashvili, Yu-Seng Huang, Ching-Yi Huang, Kim-Teng Lee, Yi-Chieh Wang, Yu Wu, Hao-Xun Chang, Chih-Horng Kuo, Erh-Min Lai, Chih-Hang Wu

## Abstract

Agroinfiltration, a method utilizing agrobacteria to transfer DNA into plant cells, is widely used for transient gene expression in plants. Besides the commonly used *Agrobacterium* strains, *Rhizobium rhizogenes* can also introduce foreign DNA into host plants for gene expression. While many *R. rhizogenes* strains have been known for inducing hairy root symptoms, their use for transient expression has not been fully explored. Here, we showed that *R. rhizogenes* A4 outperformed all other tested agrobacterial strains in agroinfiltration experiments on leaves of *Nicotiana benthamiana* and other solanaceous plants. By conducting an agroinfiltration screening in *N. benthamiana* leaves using various agrobacterial strains carrying the RUBY reporter gene cassette, we discovered that A4 mediates the strongest and fastest transient expression. Utilizing the genomic information, we developed a collection of disarmed and modified strains derived from A4. By performing vacuum infiltration assays, we demonstrated that these A4-derived strains efficiently transiently transform 6-week-old *N. benthamiana* leaves, showing less sensitivity to the age of plants compared to the laboratory strain GV3101. Furthermore, we performed agroinfiltration using AS109, an A4-derived disarmed strain, on the leaves of tomato, pepper, and eggplant. Remarkably, AS109 mediated transient gene expression on tested solanaceous plants more effectively than all the tested commonly used agrobacterial strains. This discovery paves the way for establishing *R. rhizogenes* A4-derived strains as a new option for enhancing transient expression in *N. benthamiana* and facilitating the functional study of plant genes in other solanaceous species.

## Introduction

Agrobacteria are a group of bacteria consisting of plant pathogens causing crown gall and hairy root diseases in a wide range of plants (Kado, 2014; de Lajudie *et al*., 2019; Nester, 2015). These disease symptoms arise from interkingdom gene transfer between agrobacteria and their host plants, which has driven significant research and led to the development of methods for plant transformation (Brown *et al*., 2023). This gene transfer is made possible by oncogenic plasmids within the agrobacteria genome. These plasmids, which carry most of the genetic information required for pathogenicity, are classified as tumor-inducing plasmids (pTi), responsible for crown galls, or root-inducing plasmids (pRi), which cause hairy roots (Weisberg *et al*., 2021, 2023). Together, these oncogenic plasmids provide agrobacteria with a distinctive mechanism of pathogenicity.

Agrobacteria are classified into three biovars, each representing distinct phylogenetic groups (Weisberg *et al*., 2020, 2023). Biovar 1 (BV1), traditionally known as *Agrobacterium tumefaciens*, is subdivided into several genomospecies. Among these, *A. fabacearum* (*A. tumefaciens* genomospecies 1 or G1) and *A. fabrum* (G8) have been extensively studied for their role in crown gall formation, while knowledge of the other genomospecies remains limited (Delamuta *et al*., 2020; Lassalle *et al*., 2011). Biovar 2 (BV2), historically called *Agrobacterium rhizogenes* due to its ability to induce hairy roots, has been reclassified based on genetic analyses into the genus *Rhizobium*, now known as *R. rhizogenes* (Lindström and Young, 2011; Weisberg *et al*., 2020). Biovar 3 (BV3), formerly referred to as *Agrobacterium vitis*, includes *Allorhizobium vitis* and *Allorhizobium ampelinum*, which are typically isolated from grapevines and are suspected of having a narrow host range (Kuzmanović *et al*., 2022; Mousavi *et al*., 2014, 2015). Additionally, related groups with traits similar to the three biovars have been identified, such as the BV1-like group, which includes *Agrobacterium larrymoorei* (Mousavi *et al*., 2015).

All agrobacteria sequenced to date have multipartite genomes (Weisberg *et al*., 2023). Most strains possess a primary chromosome and a circular or linear secondary chromosome, known as a chromid. While chromids exhibit plasmid-like characteristics, they also contain essential genes necessary for bacterial development (Harrison *et al*., 2010). A common characteristic of agrobacteria is the presence of oncogenic plasmids classified as pTi or pRi. These plasmids contain a transfer DNA (T-DNA) region that becomes integrated into the plant genome, and a virulence (*vir*) region which contains genes encoding the type IV secretion system (T4SS) components responsible for T-DNA delivery (Weisberg *et al*., 2023). Interestingly, the key differences between pTi and pRi is that pRi lacks *virE*, which is essential for T-DNA delivery in the pTi. Instead, pRi harbors *GALLS* which accomplish the same function as *virE* (Hodges *et al*., 2009). In addition to the oncogenic plasmids, some strains might also contain one or more non-oncogenic plasmids, whose roles in bacteria survival or pathogenicity remain largely unclear (Weisberg *et al*., 2021).

Knowledge regarding agrobacteria has been obtained using a few well-studied strains. *A. fabrum* C58 is the best-characterized strain, from which much information has been obtained (Brown *et al*., 2023; Hwang *et al*., 2017; Weisberg *et al*., 2023). Consequently, C58 has been used as a progenitor strain to generate a plethora of laboratory strains to carry out *Agrobacterium*-mediated transformation assays, including the popular strains GV3101, EHA105, AGL1 and C58C1 (De Saeger *et al*., 2021; Hwang *et al*., 2017). In addition to C58, *A. tumefaciens* Ach5 has also been previously studied, from which the laboratory strain LBA4404 was obtained (Hoekema *et al*., 1983; Ooms *et al*., 1981). Generating these strains often requires a disarming step, a process that removes the T-DNA within the oncogenic plasmid. This process eliminates the appearance of oncogenic-related symptoms in the plant tissue after infection.

Historically, plant transient transformation has been established by using disarmed, highly efficient BV1 strains (De Saeger *et al*., 2021; Hwang *et al*., 2017). In contrast, *R. rhizogenes* has traditionally been of interest for its ability to induce hairy root formation in plants (Mehrotra *et al*., 2015; Ron *et al*., 2014; Veena and Taylor, 2007). Several strains have been identified and utilized in laboratory conditions to produce hairy roots in target plant species, primarily to obtain tailor-made metabolites (Gutierrez-Valdes *et al*., 2020; Porter and Flores, 1991). Interestingly, *R. rhizogenes* is evolutionarily distant from *Agrobacterium* and *Allorhizobium* (Weisberg *et al*., 2023). Although *R. rhizogenes* and *Agrobacterium* have diverged evolutionarily, they still share similar mechanisms for transferring DNA to plants via their oncogenic plasmids (Weisberg *et al*., 2023). However, their chromosomal genetic repertoires can differ substantially. For example, certain *R. rhizogenes* genomes harbor a putative type III secretion system (T3SS) gene cluster, a nanomachine used by some rhizobia for nodulation, which is absent in other agrobacterial strains (Tampakaki, 2014). These distinctions suggest that rhizogenic BV2 strains may use a distinct plant infection strategy, potentially influencing their effectiveness in plant transformation.

One of the most common uses of agrobacteria is the transient expression of specific genes in plant leaves through a technique known as agroinfiltration (Vaghchhipawala *et al*., 2011). This method involves using a needleless syringe to introduce the bacteria, which is transformed with a binary vector containing the gene of interest, into the extracellular leaf spaces. The gene of interest within the T-DNA borders is then transferred into and expressed in the plant cells. This technique is typically applied to a limited number of plant species, with the solanaceous plant *Nicotiana benthamiana* being the most favored due to its ease of agroinfiltration and the rapid, clear gene expression observed afterward (Bally *et al*., 2018).

The transient expression system in *N. benthamiana* has been employed in various applications, including cell death assays, cell biology studies, metabolite production, molecular pharming, and recombinant protein synthesis (Bally *et al*., 2018; Goodin *et al*., 2008). These diverse applications have driven efforts within the scientific community to enhance and optimize transient expression efficiency. Recent efforts have focused on adapting plant hosts to improve gene expression efficiency (Beritza *et al*., 2024), alongside advancements in bacterial strains used for plant transformation (De Saeger *et al*., 2021). Another common strategy involves the development of adjuvant plasmids for virulence, known as ternary vectors, which typically carry additional copies of *vir* genes or genes that enhance bacterial colonization and T-DNA delivery (Aliu *et al*., 2024; Jeong *et al*., 2024). Other approaches aim to boost transformation efficiency by introducing novel elements, such as the T3SS from *Pseudomonas syringae* (Raman *et al*., 2022). Overall, most improvements have focused on refining the already well-adapted laboratory strains, specifically the BV1 strains. However, the creation of new laboratory strains from different genetic backgrounds remains a largely untapped area of research.

In this study, we identified *R. rhizogenes* A4 as the top-performing agrobacterial strain among 47 evaluated using agroinfiltration in *N. benthamiana*. We confirmed this by comparing it with several laboratory strains, measuring betalain production, GFP fluorescence, and cell death intensity. To enhance its utility, we developed several A4-derived strains, all of which retained the original transformation efficiency. Using AS109, an A4-derived disarmed strain, we demonstrated successful transient expression in 6-week-old *N. benthamiana* and superior reporter expression in tomato, pepper, and eggplant compared to other laboratory agrobacterial strains. These results emphasize the potential of A4-derived strains for transient expression in *N. benthamiana* and functional studies in other solanaceous plants.

## Results

### *R. rhizogenes* A4 is more efficient than several agrobacteria in transient expression in *N. benthamiana*

Agrobacteria, a diverse group of bacteria capable of T-DNA delivery, have become essential tools in research. While the commonly used laboratory strains are often derived from a few wild strains such as C58 and Ach5, the potential of other wild strains with distinct genetic backgrounds for robust transient gene expression remains largely underexplored. To investigate this, we performed an agroinfiltration screening in *N. benthamiana* using 47 agrobacterial strains, encompassing several BV1 strains alongside strains from the BV1-like, BV2, and BV3 groups with diverse genetic backgrounds. (Figure 1). We transformed these strains with a plasmid containing the RUBY reporter under the control of a CaMV 35S promoter for in-planta expression. The RUBY reporter system facilitates the conversion of tyrosine into betalain, resulting in visible purple pigmentation that can be easily quantified (He *et al*., 2020). At 2 days post-infiltration (dpi), leaf discs were collected, and betalain content was measured as an indicator of transient expression efficiency. The results showed significant variation in expression efficiency across different biovars and genomospecies (Figure 1).

**Figure 1.**
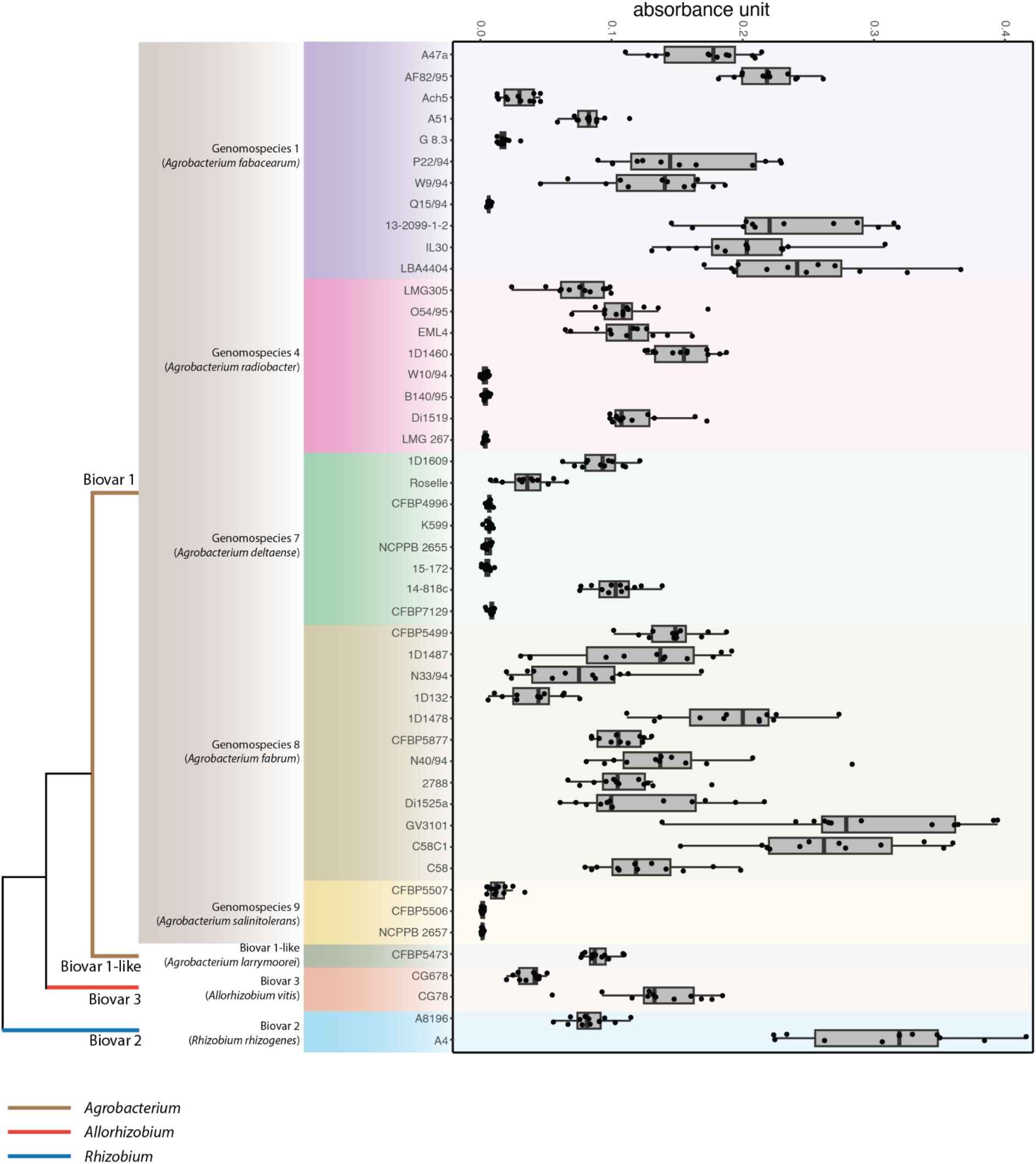
*R. rhizogenes* A4 stands out as one of the top-performing strains among 47 agrobacterial strains tested for transient expression in *N. benthamiana* leaves. Left: simplified phylogenetic tree of the four agrobacteria biovars included in this screening. Each branch color indicates different genera, and the species and biovars are indicated. Right: betalain accumulation assays using 47 agrobacterial strains (n = 12). In box plots, the line inside the box indicates the median, the edges of the box represent the 25th and 75th percentiles, and the whiskers reach out to the most extreme data points, up to 1.5 times the interquartile range. Different color backgrounds indicate different species.

Among the BV1 strains, G1 and G8 strains demonstrated higher betalain accumulation overall compared to those belonging to G4, G7, and G9 (Figure 1). Many of the top-performing BV1 strains were already well-established laboratory strains, including LBA4404 and GV3101 from G1 and G8, respectively. For the non-BV1 strains tested here, only one BV2 strain, A4, exhibited high betalain accumulation (Figure 1). This strain, which has been used in producing hairy roots for metabolite production (Gharari *et al*., 2020; Królicka *et al*., 2001; Petit *et al*., 1983; Sathasivam *et al*., 2022), unexpectedly exhibited transient expression efficiency similar to GV3101, C58C1, and LBA4404, three of the most commonly used laboratory strains (Figure 1).

To further compare A4 with the strains showing the highest transient expression efficiency, we performed side-by-side comparisons using several strains on the same *N. benthamiana* leaves. These included GV3101, a widely used strain for agroinfiltration in this species, C58C1, known for its high efficiency in transient gene expression in *Arabidopsis* seedlings using the AGROBEST method, LBA4404, a strain commonly used for stable transformation in other solanaceous species, and Ach5, the G1 strain from which LBA4404 was derived (De Saeger *et al*., 2021; Hwang *et al*., 2017; Wu *et al*., 2014) (Figure 2a,b). Surprisingly, the leaf spots infiltrated with A4 exhibited the highest betalain accumulation compared to all other strains at 18, 24, and 48 hours post-infiltration (hpi) (Figure 2a,b). Notably, pigmentation appeared more rapidly in A4-infiltrated areas, with visible coloration as early as 18 hpi.

**Figure 2.**
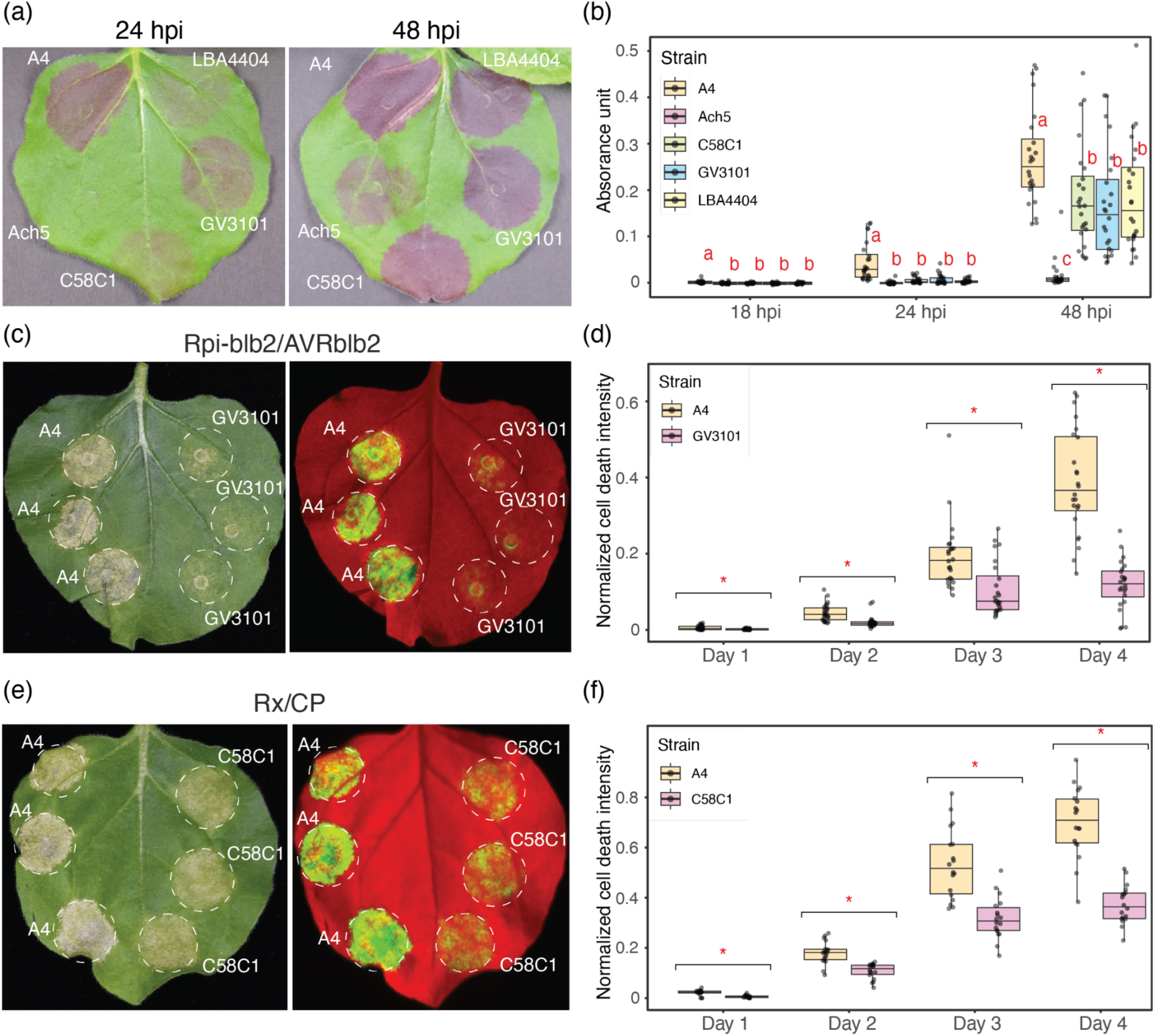
*R. rhizogenes* A4 is more efficient than several commonly used laboratory agrobacterial strains in transient expression in *N. benthamiana.* (a) Evaluation of transient expression efficiency of A4, LBA4404, Ach5, GV3101 and C58C1 agrobacterial strains on the same *N. benthamiana* leaf using the RUBY reporter. Photos were taken at 24 and 48 hours post-infiltration (hpi). (b) Betalain absorbance measurements were performed on the infiltration spots shown in (a) at three different timepoints (n = 24). Letters indicate statistically significant differences between treatments at each timepoint (Tukey’s HSD test, p < 0.05). (c-f) Cell death assays comparing A4 with GV3101 or C58C1, using two different effector/receptor pairs. Photos were taken at 4 days after co-infiltration with Rpi-blb2 and AVRblb2 (c) or Rx and CP (e). Quantification of the cell death assays for Rpi-blb2/AVRblb2 (d) and Rx/CP (f) was performed at four different timepoints (n = 12). In box plots, the line inside the box indicates the median, the edges of the box represent the 25th and 75th percentiles, and the whiskers reach out to the most extreme data points, up to 1.5 times the interquartile range. * indicate statistically significant differences between treatments at each timepoint (Dunn’s test, p < 0.05).

We next compared A4 with another two commonly used laboratory BV1 strains, AGL1 and EHA105, which were not included in our initial screening. The result demonstrated that A4 outperformed these strains on *N. benthamiana* leaves (Figure S1). Additionally, we included the rhizogenic strain ARqua1, which carries the C58 chromosomal background with pRiA4b, and A8196, another BV2 strain that harbors a type II pTi, to assess the effects of plasmid and chromosomal backgrounds (Figure S2). Our results indicated that A4 outperformed both of these strains. These results also suggest that the high efficiency of A4 cannot be simply explained by its oncogenic plasmid type or chromosomal background differences between biovars.

To validate the results obtained with the RUBY reporter, we conducted an agroinfiltration assay using GFP expressed under a 35S promoter for constitutive in-planta expression. GFP fluorescence quantification confirmed faster and stronger fluorescence at A4 infiltration sites compared to GV3101 (Figure S3a,b). Moreover, GFP protein levels were significantly higher with A4, mirroring the fluorescence patterns observed (Figure S3c).

We then performed hypersensitive cell death assays by co-expressing pathogen avirulence factors and their corresponding plant immune receptors (Figure 2c-f). We co-expressed AVRblb2, an effector from the oomycete *Phytophthora infestans*, with the corresponding immune receptor Rpi-blb2 from *Solanum bulbocastanum* (Oh *et al*., 2014). Cell death occurred earlier at the A4 infiltration sites compared to GV3101, and the signal remained stronger up to 4 dpi (Figure 2c,d). A similar experiment, expressing the coat protein from Potato Virus X alongside its immune receptor Rx from *Solanum tuberosum* (Bendahmane *et al*., 1995), produced consistent results. Cell death phenotypes appeared earlier and were more pronounced at the A4 infiltration sites compared to C58C1 (Figure 2e,f). Overall, A4 consistently proved to be the most effective strain for transient transformation in *N. benthamiana*, surpassing all other strains tested in our assays.

### Modifications of A4 to generate laboratory strains

A4 is typically used for generating hairy roots for metabolite production but has not been employed for transient expression in leaves. During our transient expression experiments with A4, we observed leaf curling starting two days after bacterial infiltration, likely due to the expression of oncogenic T-DNA genes, which is undesirable for such applications (Figure S4a). Additionally, when untransformed A4 was plated on kanamycin (50 mg/L), a few kanamycin-resistant colonies emerged spontaneously (Figure S4b). Consequently, we decided to modify A4 to generate more user-friendly strains for routine laboratory use.

To achieve this, we sequenced the complete genome of A4, identifying four replicons: a circular chromosome (chromosome 1), a circular chromid (chromosome 2), a Ri plasmid (pRiA4), and a non-oncogenic plasmid (pArA4) (Figure 3a). Gene annotation enabled us to locate both the *vir* region and the two T-DNA regions within pRiA4. We also discovered a putative kanamycin resistance gene on chromosome 1, which may account for the spontaneous kanamycin-resistant phenotype observed (Figure 3a and Figure S4b).

**Figure 3.**
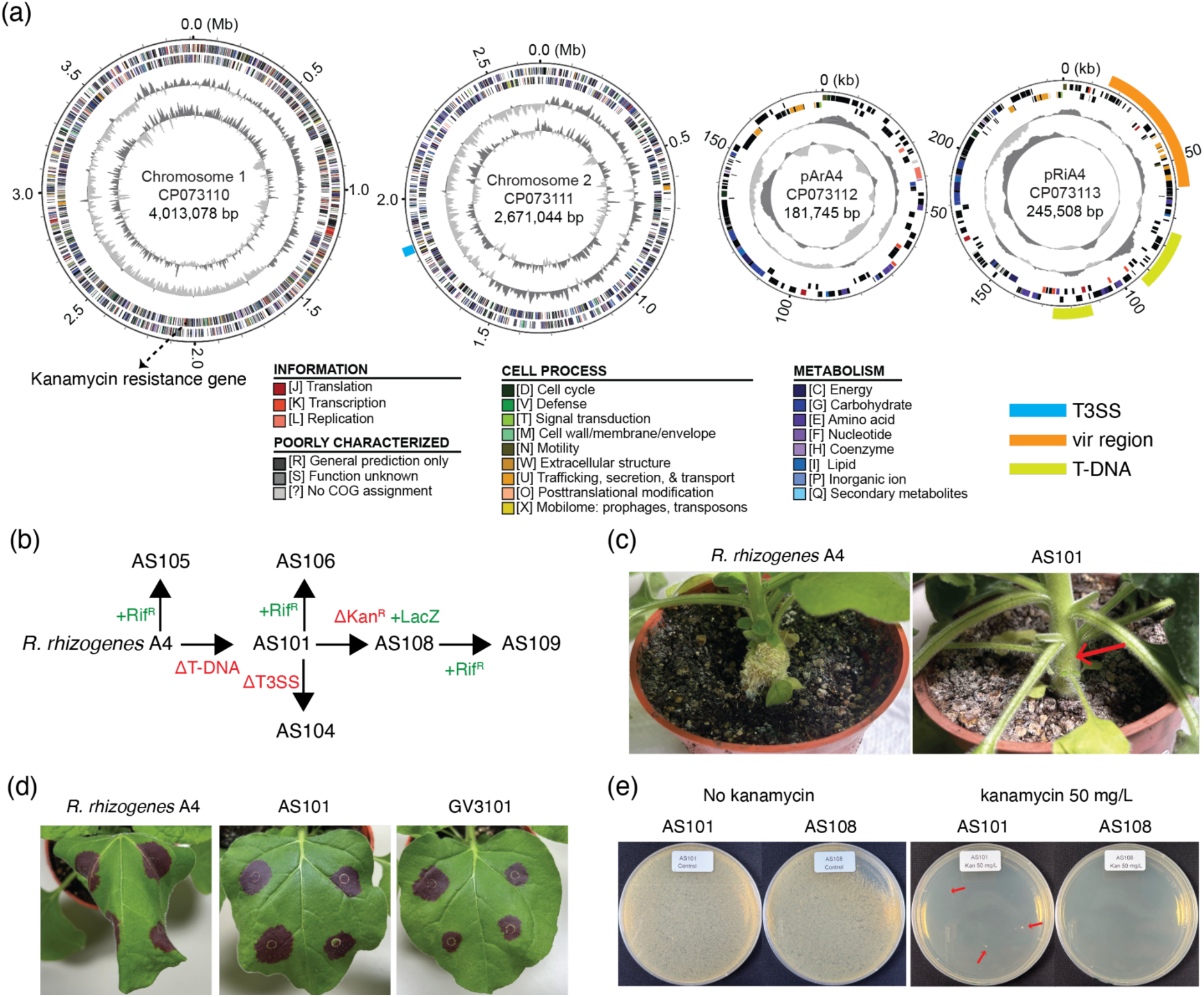
Genomic information of *R. rhizogenes* A4 and modification of A4 for routine use in the laboratory. (a) Genome map of *R. rhizogenes* A4. The rings, from outermost to innermost, represent: (1) scale marks; (2) coding sequences on the forward strand; (3) coding sequences on the reverse strand; (4) GC skew (dark or light gray color indicates positive or negative values, respectively); (5) GC content (dark or light gray color indicates above or below average values, respectively). Putative functions of identified coding sequences and the type III secretion system (T3SS), T-DNA, and *vir* region are indicated. (b) Schematic representation of the modifications made to *R. rhizogenes* A4 to generate A4-derived strains. Green text indicates additions to the genome, while red indicates deletions. (c) Hairy root induction assay comparing *R. rhizogenes* A4 and AS101. No hairy roots were observed at the AS101 infection site (arrow). (d) Comparison of leaf curling phenotypes between *R. rhizogenes* A4, the disarmed strain AS101 and GV3101. Pictures were taken 48 hours post-infiltration. (e) Kanamycin susceptibility test comparing AS101 and AS108. Antibiotic-resistant mutants of AS101 appeared on 523 medium plates containing kanamycin 50 mg/L (arrows), while no AS108 colonies were observed.

Using a homologous recombination-based approach, we successfully removed both T-DNA regions, resulting in a strain designated as AS101 (Figure 3b). To confirm the success of this disarming process, we conducted a hairy root production assay on *N. benthamiana* stems using both A4 and AS101, monitoring hairy root formation three weeks post-inoculation (Figure 3c). While A4 induced ectopic root formation on the stems, AS101 did not exhibit this phenotype, confirming the success of the disarming (Figure 3c). We then carried out transient expression experiments on *N. benthamiana* leaves, comparing A4, AS101, and GV3101. As anticipated, A4 induced the leaf curling phenotype previously described, whereas the disarmed AS101 and the laboratory strain GV3101 did not cause any leaf curling (Figure 3d).

Using AS101 as the genetic background and the INTEGRATE technology (Aliu *et al*., 2024; Vo *et al*., 2021), we disrupted the endogenous, putative kanamycin-resistance gene. To avoid interference with future cell biology experiments, we modified the plasmid pEA186 by replacing the Tn cargo mCherry cassette with a LacZ cassette. We subsequently generated another strain, AS108, by disrupting the putative kanamycin resistance gene with a LacZ cassette. To evaluate the kanamycin sensitivity of AS108, we conducted a kanamycin sensitivity assay, comparing both A4 and AS108 (Figure 3e and Figure S5a). As expected, A4 exhibited the spontaneous emergence of kanamycin-resistant colonies within just 2 days. In contrast, AS108 failed to produce any colonies at the tested kanamycin concentrations, confirming that the targeted gene was responsible for the sporadic development of resistance (Figure S5a).

Since most commonly used laboratory strains exhibit resistance to rifampicin, we introduced this trait into A4, AS101, and AS108 by exposing them to serial concentrations of the antibiotic, resulting in the creation of AS105, AS106, and AS109, respectively. We confirmed that A4, AS101, and AS108 were unable to grow on media containing rifampicin at 50 mg/L, while AS105, AS106, and AS109 successfully grew under the same conditions (Figure S5b).

### A4-derived strains maintain hyper-efficient transient expression efficiency on *N. benthamiana*

To verify that the newly generated A4-derived strains retained their hyper-efficient transient expression capability in *N. benthamiana*, we conducted an agroinfiltration comparison using the RUBY reporter for strains A4, AS101, AS105, AS106, AS108, and AS109, infiltrating them into the same leaf (Figure 4a,b). Betalain accumulation showed no significant differences among the six strains at 18, 24, and 48 hpi, confirming that the hyper-efficient phenotype remained unchanged on these laboratory strains.

**Figure 4.**
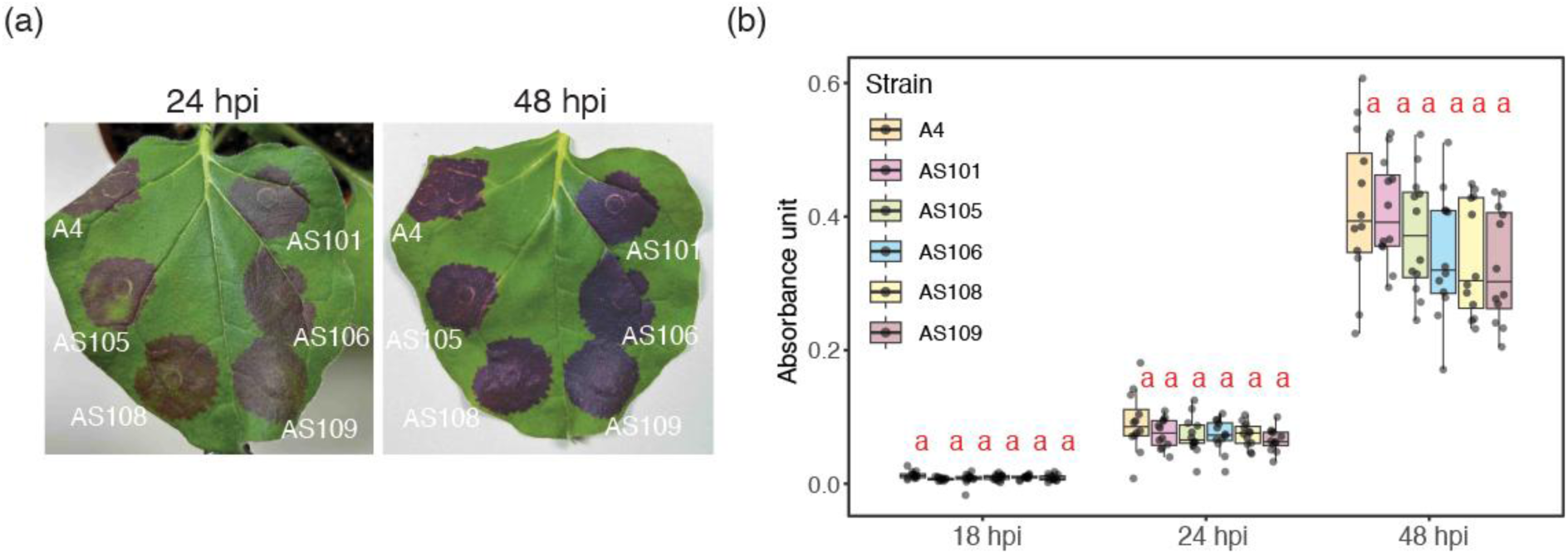
Strains derived from *R. rhizogenes* A4 consistently exhibit highly efficient transient expression in *N. benthamiana*. (a) Evaluation of transient expression efficiency of A4 and the A4-derived strains AS101, AS105, AS106, AS108 and AS109 on the same *N. benthamiana* leaf using the RUBY reporter. Photos were taken at 24 and 48 hours post-infiltration. (b) Betalain absorbance measurements were performed on the infiltration spots shown in (a) at three different timepoints (n = 12). In box plots, the line inside the box indicates the median, the edges of the box represent the 25th and 75th percentiles, and the whiskers reach out to the most extreme data points, up to 1.5 times the interquartile range. Letters indicate statistically significant differences between treatments at each timepoint (Tukey’s HSD test, p < 0.05).

### A4-derived strains mediate transient expression on *N. benthamiana* in a less age-sensitive way

Previous studies showed that *Agrobacterium*-mediated transient expression in *N. benthamiana* is sensitive to the age of the plant, with older plants being more recalcitrant to transient transformation (Dodds *et al*., 2023; Saur *et al*., 2016). To test whether this is also the case for *R. rhizogenes* A4-derived strains, we performed vacuum infiltration of A4, AS109, and GV3101 on both 4-week-old and 6-week-old plants. Three days after the vacuum infiltration, we quantified betalain accumulation in the 5th to the 11th true leaves for 6-week-old plants. In the 4-week-old plants, A4, AS109 and GV3101 mediated efficient RUBY reporter expression, with no obvious differences at 72 hpi (Figure S6). In the 6-week-old plants, GV3101 showed very low transient expression efficiency, with the upper (10th and 11th) leaves showing slightly higher betalain accumulation compared to the lower leaves (Figure 5). Interestingly, both A4 and AS109 mediated clear betalain accumulation from the lower to the upper leaves, with the upper leaves showing slightly stronger betalain accumulation (Figure 5). Across all leaves in 6-week-old plants, A4 and AS109 mediated stronger betalain accumulation compared to GV3101 (Figure 5).

**Figure 5.**
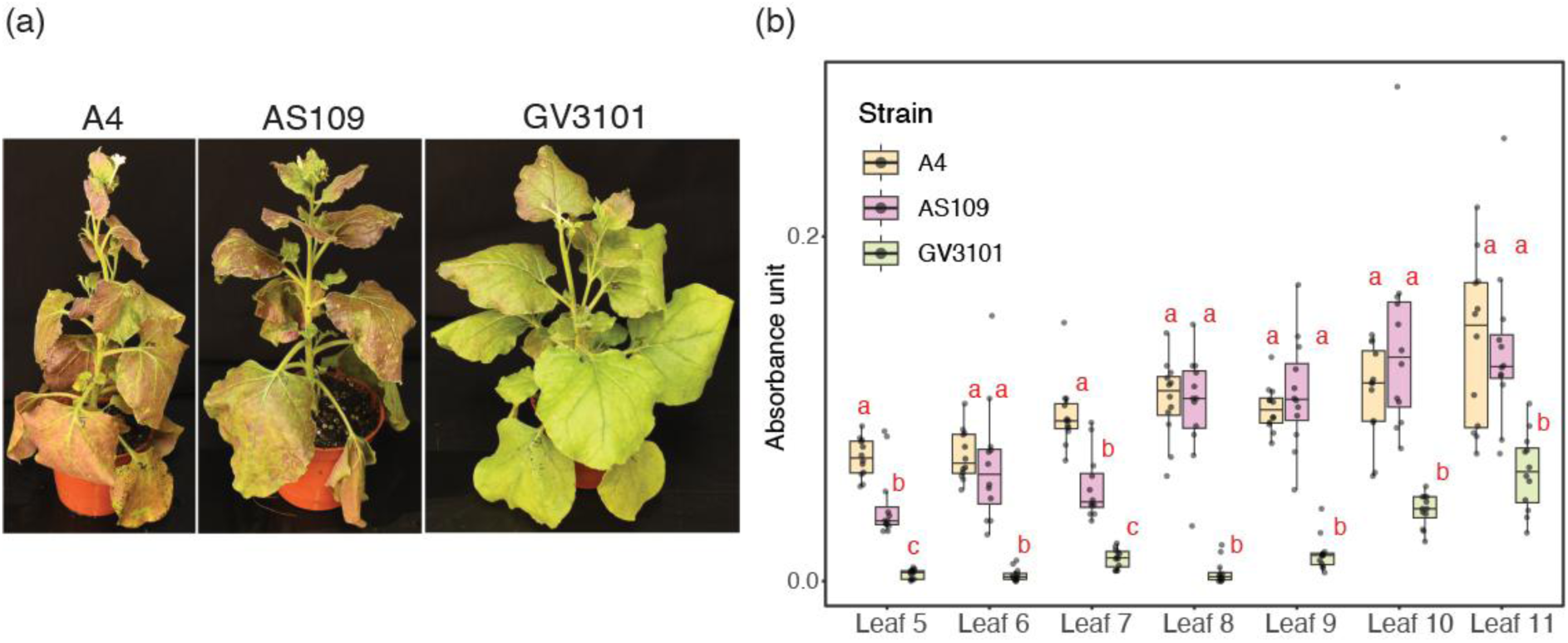
*R. rhizogenes* A4-derived strains exhibit less sensitivity to the age of *N. benthamiana* in transient expression experiments. (a) Transient expression efficiency on 6-week-old *N. benthamiana* leaves of A4, AS109 and GV3101 using vacuum infiltration and the RUBY reporter. Photos were taken at 3 days post-infiltration. (b) Betalain absorbance measurements were performed on the 5th to the 11th leaf (counting from the bottom), by collecting leaf discs from three different plants (n = 12). In box plots, the line inside the box indicates the median, the edges of the box represent the 25th and 75th percentiles, and the whiskers reach out to the most extreme data points, up to 1.5 times the interquartile range. Letters indicate statistically significant differences between treatments at each timepoint (Tukey’s HSD test, p < 0.05).

### A4-derived strains outperformed other laboratory strains on transient expression in leaves of solanaceous plants

Although agroinfiltration in *N. benthamiana* has been widely employed, its use in other solanaceous plants has been limited. To evaluate the suitability of *R. rhizogenes* A4-derived strains for transient expression in other solanaceous plants, we conducted agroinfiltration experiments comparing AS109 with various laboratory strains in the leaves of eggplant, tomato, and pepper—three economically significant solanaceous crops. In eggplant (cv. HV-037) leaves, AS109, C58C1, EHA105, and LBA4404 outperformed AGL1 and GV3101, with AS109 yielding the highest average betalain accumulation (Figure 6a,b). In both tomato (cv. Moneymaker) and pepper (cv. Groupzest), AS109 was the only strain to produce notable betalain accumulation at 2 dpi (Figure 6a,b). We then explored the potential of AS109 for cell death assays in these plants. Using AS109 and C58C1, we transiently expressed Rx and CP in eggplant, tomato, and pepper leaves. While Rx and CP expression via C58C1 induced minimal or no visible cell death across the tested species, expression with AS109 consistently triggered cell death in all cases (Figure 6c-e). No visible symptoms appeared when Rx and CP were expressed individually using AS109 or C58C1. These results suggest that AS109 outperforms the tested agrobacterial strains for transient expression in eggplant, tomato, and pepper and holds promise for further functional genomics studies in these crops.

**Figure 6.**
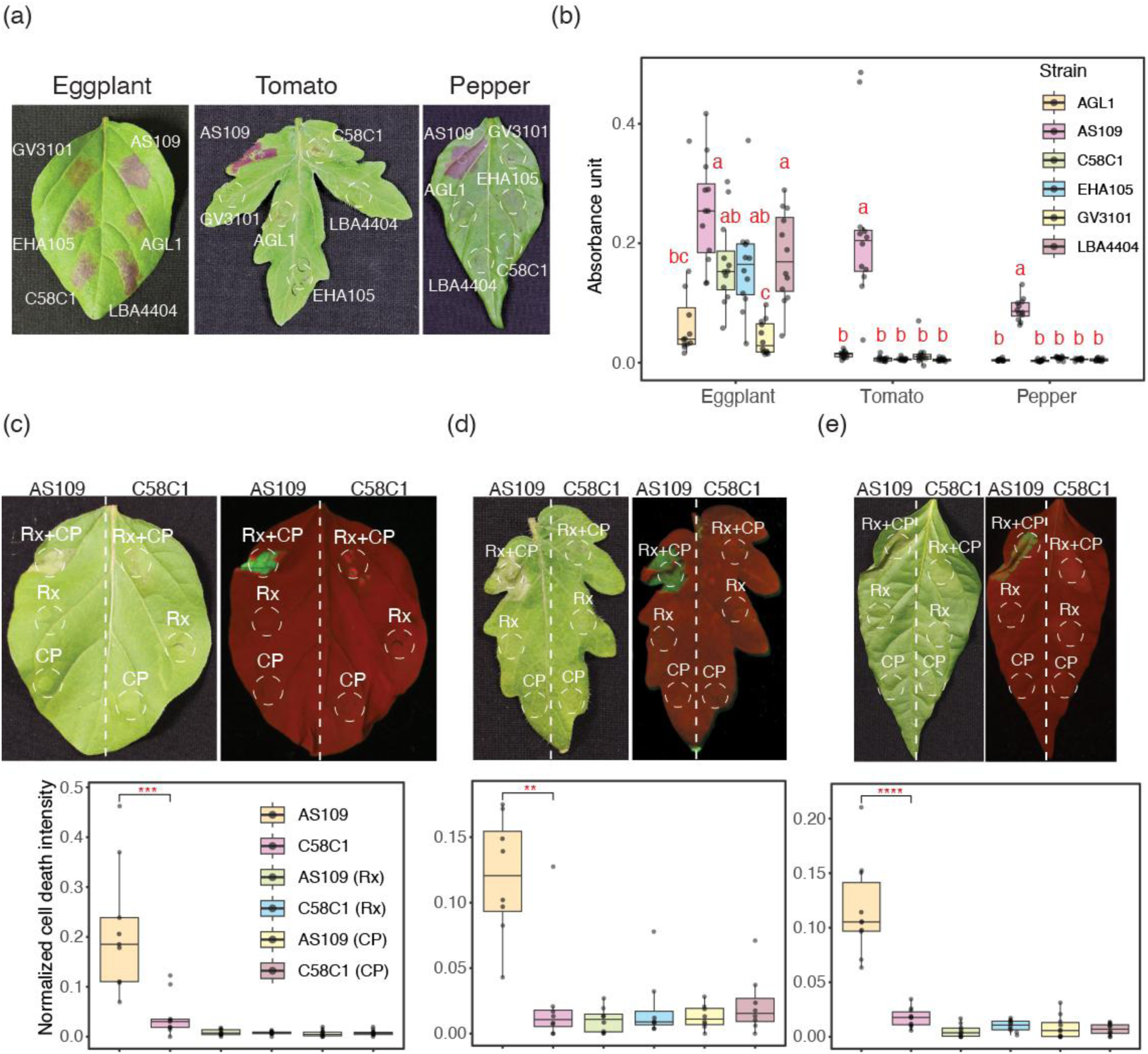
*R. rhizogenes* A4-derived strains AS109 outperformed other laboratory strains on transient expression in tomato, pepper, and eggplant leaves. (a) Evaluation of transient expression efficiency of AS109, GV3101, EHA105, AGL1, C58C1 and LBA4404 agrobacterial strains on the same eggplant (cv. HV-037), tomato (cv. Moneymaker) or pepper (cv. Groupzest) leaf using the RUBY reporter. Photos were taken at 48 hours post-infiltration. (b) Betalain absorbance measurements were performed on the infiltration spots shown in (a) (n = 12). Letters indicate statistically significant differences between treatments at each timepoint (Tukey’s HSD test, p < 0.05). (c-e) Cell death assays comparing A4 with C58C1, using CP and Rx effector/receptor pairs on eggplant (cv. HV-037) (c), tomato (cv. Moneymaker) (d) or pepper (cv. Groupzest) (e) leaves. Photos were taken and quantification was performed at 2 days after co-infiltration (n = 9 in eggplant; n = 10 in pepper; n = 8 in tomato). * indicate statistically significant differences between treatments at each timepoint (Wilcoxon signed rank test, ** = p < 0.01, *** = p < 0.001, **** = p < 0.0001). In box plots, the line inside the box indicates the median, the edges of the box represent the 25th and 75th percentiles, and the whiskers reach out to the most extreme data points, up to 1.5 times the interquartile range.

## Discussion

Through a comprehensive screening of various agrobacterial strains in *N. benthamiana* leaves using a RUBY reporter gene cassette, we identified *R. rhizogenes* A4 as the top-performing strain, demonstrating the fastest and most robust betalain accumulation. Additionally, A4-derived strains proved effective for transiently transforming other solanaceous plants and 6-week-old *N. benthamiana* plants, in contrast to the limited efficiency observed with other laboratory strains. These findings highlight A4-derived strains as valuable tools for enhancing transient expression in *N. benthamiana* and for advancing functional gene analysis in other solanaceous species.

Agrobacteria have long been valuable tools for plant biotechnology, particularly for transient gene expression, which allows for rapid and temporary expression of foreign genes in plants. To this end, different agrobacterial strains, such as *A. fabrum* (e.g., GV3101, EHA105) and *A. fabacearum* (e.g., LBA4404), have been developed and optimized for specific applications. Early studies on transient expression identified strains that could be used in particular plant species (Hwang *et al*., 2013; Wroblewski *et al*., 2005). Similar to transient gene expression, several strains have been tested in early research to establish stable transformation protocols (Hwang *et al*., 2017). Interestingly, despite sharing the same chromosomal background, closely related agrobacterial strains often exhibit drastically different performances across plant species.

Significant progress has been made over the last four decades in understanding agrobacterial biology, including the identification of pTi *vir* genes, chromosomal virulence (*chv*) genes, the discovery of T4SS for T-DNA transfer and type VI secretion system for niche competition (Brown *et al*., 2023; Weisberg *et al*., 2023). However, much of this research has centered on *A. fabrum* and *A. fabacearum* strains within BV1 (De Saeger *et al*., 2021). This narrow focus has left a significant gap in our understanding and utilization of non-BV1 strains, creating a bottleneck in agrobacteria research. Our study demonstrates that the BV2 strain *R. rhizogenes* A4 outperforms traditionally used BV1 *Agrobacterium* strains in several solanaceous plants, indicating the untapped potential within the natural diversity of agrobacterial resources. By exploring and engineering the genetic backgrounds of diverse agrobacterial strains, it is possible to develop more efficient and versatile expression systems, providing tailored tools for various plant species and advancing biotechnology applications.

Although transient expression in pepper (Liu *et al*., 2024; Oh *et al*., 2023; Zhang *et al*., 2020) and tomato (Hoshikawa *et al*., 2019) has been previously investigated, most studies rely on C58-derived strains. While we did not directly test the protocols outlined in these studies, our experience with C58-derived strains in pepper and tomato yielded unsatisfactory results. Alternatively, some researchers used viral vectors to enhance protein expression (Hoshikawa *et al*., 2019; Liu *et al*., 2024), but these methods may not be suitable for some experimental designs. Our study found that A4-derived strains, such as AS109, outperformed other strains in transient expression across pepper, tomato, and eggplant, with no observable side effects. Additionally, we consistently observed expected phenotypes, including RUBY expression and cell death, in all leaves tested for these crops, demonstrating the reliability of AS109 in these assays. The infiltration protocol for AS109 aligns closely with the methods used in most of our other infiltration experiments, making it straightforward for other laboratories to adapt. The broad host range of A4 strains within the Solanaceae family highlights their potential application in economically important species. However, their utility beyond the Solanaceae remains unexplored. Future research will aim to uncover the full potential of A4 strains as a versatile tool for transient expression across a wider range of species.

Recent efforts to enhance plant transformation efficiency have largely focused on manipulating plant host responses to increase receptivity to genetic modifications. A recent review by Beritza et al. (2024) underscores the role of subcellular processes that impede transient expression in *N. benthamiana*. Strategies to mitigate these issues include delivering effectors that suppress plant immune responses, preventing proteolysis of accumulated proteins, or targeting the expression of proteins to specific subcellular compartments (Beritza *et al*., 2024). Another promising approach involves the development of new plasmid systems, such as ternary vectors, which can express additional copies or mutants of virulence genes (Aliu *et al*., 2024; Jeong *et al*., 2024). These vectors may also include genes that suppress or compromise the plant immune system, thereby enhancing gene transfer efficiency. It would be valuable to explore how these methods could be integrated with the use of A4-derived strains to further maximize transient expression efficiency in *N. benthamiana*. Combining these complementary approaches may provide a robust framework for advancing plant genetic engineering techniques.

Exploring the mechanisms behind the higher efficiency of *R. rhizogenes* A4 compared to commonly used laboratory strains remains an intriguing area of investigation. However, our initial attempts to identify the underlying factors were unsuccessful. Our first hypothesis was that effector protein(s), potentially secreted via the T3SS, interferes with plant immune responses. This hypothesis was based on the absence of the T3SS in BV1 strains and previous findings showing enhanced transformation efficiency in *A. fabrum* strain EHA105 engineered to express T3SS and effectors from *Pseudomonas syringae* (Raman *et al*., 2022). However, knocking out the putative T3SS gene cluster in A4 did not affect its efficiency in *N. benthamiana* (Figure S7). This result suggested that either A4 does not produce a functional T3SS or the T3SS effectors are not a significant factor in the hyper-efficiency of A4. Another intriguing characteristic of A4 is its ability to transiently transform 6-week-old *N. benthamiana* plants, a feature not achieved by GV3101. This bypass of age-dependent resistance in *N. benthamiana* was hypothesized to be related to the csp22 epitope, a known trigger of plant immune responses induced by agrobacterial cold shock proteins (Dodds *et al*., 2023; Saur *et al*., 2016). We speculated that A4 might have csp22 epitopes too divergent to be recognized by plant receptors, allowing it to evade immune detection. However, our analysis revealed that csp22 epitopes of A4 are similar to those of GV3101 (Figure S8). Whether these epitopes differ in their ability to trigger immune responses or whether the cold shock proteins are actively expressed during infection remains an open question. Future studies employing RNA sequencing to profile transcriptional changes during infection or performing mutant screens on A4 could help identify the key factors contributing to its superior efficiency. These approaches may uncover the molecular mechanisms underpinning the hyper-efficient phenotype of A4, paving the way for broader applications and further optimization.

In conclusion, our exploration of agrobacterial diversity has identified *R. rhizogenes* A4 as a highly effective tool for transient expression in various solanaceous plant species. Although the applications of A4 beyond solanaceous plants remain untested, this discovery underscores the importance of exploring the diverse genetic backgrounds within this complex bacterial group. Such investigations can deepen our understanding of their virulence mechanisms and help identify strains with substantial scientific and economic potential. The exceptional hyper-efficient transient expression efficiency of A4 presents an exciting opportunity for further investigation in follow-up studies. Future work will focus on expanding its applications in solanaceous plants, including cell biology assays and molecular farming. Promising avenues such as stable transformation and gene editing in solanaceous plants will also be explored. We believe the discovery of A4, especially when combined with innovative methods to enhance transformation efficiency, such as ternary vectors, has the potential to revolutionize routine transient expression assays in the years to come.

## Material and Methods

### Plant material and growth conditions

Wild-type *N. benthamiana*, eggplant (cv. HV-037), tomato (cv. Moneymaker) and pepper (cv. Groupzest) plants were grown in a walk-in chamber with a temperature of 25°C, humidity of 45–65%, and 16/8 hr light/dark cycle. Eggplant seeds were purchased from Known-You Seed Co., Ltd., and pepper seeds were a gift from Dr. Miin-Huey Lee (National Chung Hsing University, Taiwan).

### Agrobacterial strains

All agrobacterial strains used in this study are listed in Table S1. Strains were subcultured on 523 medium at 28°C (1% sucrose, 0.8% casein hydrolysate (N-Z-Case), 0.4% yeast extract, 0.2% K_2_HPO_4_, and 0.03% MgSO_4_·7H_2_O, adjusted to pH 7) by using agar plates (containing 1.5% agar) or liquid 523 media (300 rpm shaking), unless otherwise specified.

### Agroinfiltration

Agroinfiltration experiments were conducted as described previously (Chiang *et al*., 2024). Briefly, agrobacteria carrying the appropriate binary vectors were cultured on solid 523 medium for 1–2 days, then resuspended in MMA buffer (10 mM MgCl_2_, 10 mM MES-KOH, 200 µM acetosyringone, pH 5.6). The bacterial suspension was adjusted to the appropriate optical density (OD_600_) before infiltrating the leaves of 4-week-old *N. benthamiana*. For eggplant, tomato and pepper, 3 to 4-week-old plants were used. Details on the OD_600_ values used in each assay are provided in Table S2.

### Vacuum infiltration in *N. benthamiana*

Agrobacteria for vacuum infiltration were prepared as above. To prevent soil contamination in the infiltration solution, the soil surrounding 4 or 6-week-old *Nicotiana benthamiana* plants was covered. The entire plant was then submerged in a container with the infiltration solution and placed in a vacuum chamber. A vacuum was applied using a pump at a pressure of at least 500 mBar for 1.5 minutes. After infiltration, the plants were removed from the solution and transferred to a growth chamber for 72 hours.

### RUBY quantification

RUBY quantification was carried out as previously described (Chiang *et al*., 2024; Polturak *et al*., 2017). Briefly, leaf discs were collected and incubated in an extraction buffer containing 10% ethanol and 0.1% formic acid. The quantification was then performed using a spectrophotometer.

### Cell death and GFP quantification

Cell death autofluorescence or GFP fluorescence of leaves was quantified as described previously (Goh *et al*., 2024). Briefly, raw autofluorescence or fluorescence images of the infiltration sites were captured using the UVP ChemStudio PLUS (Analytik Jena), with blue LED light for excitation and a FITC filter (513–557 nm) for emission. The mean signal intensity within the selected infiltrated area was normalized to the maximum possible intensity (65535) to calculate the normalized cell death intensity or fluorescence density.

### Protein extraction and western blot analysis

SDS-PAGE and western blot analysis were carried out as described by Chiang et al. (2024). Briefly, infiltration sites expressing GFP were collected 1 to 4 days post-infiltration (dpi) and flash-frozen in liquid nitrogen. Following protein extraction, the samples were heated at 70°C for 10 minutes prior to SDS-PAGE and immunodetection. Anti-GFP (YH80006; Yao-Hong) served as the primary antibody, while Peroxidase-Conjugated Goat Anti-Rabbit IgG (H + L) (AP132P; Sigma) was used as the secondary antibody. Images were captured using the UVP ChemStudio PLUS (Analytik Jena). SimplyBlue SafeStain (465034; Invitrogen) was employed to visualize Rubisco on the PVDF membrane.

### Genome sequencing and analysis

The procedure of genome sequencing and analysis of A4 was modified from that described in our previous study (Huang *et al*., 2015). Briefly, a random genomic DNA library was prepared and sequenced on the Illumina MiSeq platform to obtain 4,346,861 pairs of 300-bp reads, providing ~367-fold coverage of this 7.1 Mb genome sequence. The quality-trimmed reads were assembled using Velvet v1.2.10 (Zerbino and Birney, 2008) with the k-mer length set to 151. The contigs were oriented and assembled into scaffolds using the complete genome sequence of K599 (NCBI RefSeq accession GCF_002005205.2) as the reference. The gaps were filled by mapping the Illumina raw reads to the scaffolds using BWA v0.7.17 (Li and Durbin, 2009) for extending the contigs or by PCR across contigs followed by Sanger sequencing until the complete sequences of all four circular replicons were determined. For validation, all Illumina raw reads were mapped to the finalized assembly using BWA v0.7.17, followed by programmatically checks using SAMTOOLS v1.9 (Li *et al*., 2009). The annotation was provided by NCBI using the Prokaryotic Genome Annotation Pipeline (PGAP) v5.1 (Tatusova *et al*., 2016). To identify the genes of interest, including those that encode antibiotic resistance or secretion systems, all predicted proteins were used as queries to perform BLASTP v2.11.0 (Boratyn *et al*., 2013) sequence similarity searches against the NCBI non-redundant protein sequences database (Sayers *et al*., 2022), followed by keyword searches of the results. Visualization of the genome map was performed using CIRCOS v0.69-9 (Krzywinski *et al*., 2009).

### Homologous recombination for bacteria gene editing

The T-DNA regions of pRiA4 were removed using a homologous recombination strategy to generate the disarmed strain AS101. For this purpose, the suicide plasmid pJQ200KS, originally with kanamycin resistance, was modified to contain a spectinomycin resistance cassette for gene replacement experiments in A4 (Quandt and Hynes, 1993). Two DNA fragments flanking the T-DNA regions were amplified using primer sets 5’XbaI-pRiA4/3’pRiA4 and 3’PstI-pRiA4/5’pRiA4, introducing XbaI and PstI restriction sites, respectively (Table S3). The amplified fragments and the acceptor plasmid were digested with XbaI and PstI (NEB), and then ligated to construct a plasmid for T-DNA removal. This plasmid was electroporated into A4 competent cells, followed by a 2-hour recovery in liquid AB-MES (pH 5.5) minimal medium at 28°C (Wu *et al*., 2014). The bacteria were then plated on an AB-MES (pH 7) solid medium containing 200 mg/L spectinomycin.

After 5–6 days, colonies containing the transformed plasmid underwent the first homologous recombination event, and then were grown in 2 mL of liquid 523 medium supplemented with 0.25% glucose (replacing sucrose) to induce the second recombination event. After 1–2 days of incubation, serial dilutions (10^-3^ to 10^-5^) of the culture were plated onto 523 solid medium containing 5% sucrose to select colonies with plasmid removal. Successful strain disarming was verified by Sanger sequencing using the primers 5’M-pRiA4, 3’M-pRiA4, and 3’B-pRiA4 (Table S3).

### INTEGRATE system for bacteria gene editing

The putative kanamycin resistance gene and the T3SS gene cluster present in the A4 genome were deleted using the INTEGRATE system following the methods described previously (Aliu *et al*., 2022; Vo *et al*., 2021).

To create the kanamycin-susceptible mutant AS108, a new INTEGRATE plasmid, pCHW027037, was generated to deliver a LacZ cassette as a Tn cargo. First, the backbone of plasmid pEA186 (Addgene #187874, described by Aliu et al., 2022) was amplified using the primer pairs pEA186_first_GG_F/pEA186_first_GG_R and pEA186_second_GG_F/pEA186_second_GG_R. The *LacZ* cassette was amplified from plasmid pAGM9121 (Addgene #51833) using primers pEA186_LacZ_GG_F/pEA186_LacZ_GG_R (Table S3). The resulting fragments were digested with BbsI (NEB) and assembled via the Golden Gate reaction to produce the INTEGRATE acceptor plasmid pCHW027037.

Next, a 32-nucleotide spacer targeting the kanamycin resistance gene was designed using the pipeline from Vo et al., 2021 and inserted into pCHW027037 after BsaI (NEB) digestion. The plasmid was then electroporated into AS101 cells, followed by a 2-hour recovery in liquid 523 medium supplemented with 0.25% glucose (replacing sucrose) at 28 °C. The cells were subsequently plated on solid 523 medium supplemented with 0.25% glucose. After 3 days, colonies were screened via PCR and colonies containing edited and non-edited bacteria were identified. Colonies were subcultured for 2–3 generations to obtain pure populations, which were then plated on 523 solid medium containing 5% sucrose to remove the plasmid. Successful kanamycin resistance gene deletion was confirmed via Sanger sequencing using the primer CRISPR_KmR_R (Table S3).

To create the T3SS deletion mutant AS104, the INTEGRATE acceptor plasmid pKL2310 was used (Aliu *et al*., 2022). Two spacers targeting the ends of the T3SS gene cluster were designed, and a DNA fragment containing both spacers was synthesized (Table S4). The fragment was then inserted into pKL2310 by using BsaI in a Golden Gate reaction. The resulting plasmid was electroporated into AS101 cells, and colonies were screened following the same procedure described for AS108 until pure, edited colonies were obtained.

To complete the deletion of the T3SS gene cluster (now flanked by LoxP sites), the edited bacteria were transformed with plasmid pKL2315 and plated on 523 medium supplemented with 0.25% glucose (replacing sucrose) and containing 125 mg/L kanamycin. Finally, the bacteria were plated on 523 medium supplemented with 5% sucrose to select the colonies that had undergone plasmid removal. Successful deletion of the T3SS gene cluster was verified through Sanger sequencing using the primer T3SSUP_spacer_F (Table S3).

### Introduction of rifampicin resistance

The rifampicin-resistant strains AS105, AS106, and AS109 were derived from A4, AS101, and AS108, respectively. For this purpose, 5 mL cultures of each strain in liquid 523 medium containing 0.5 mg/L rifampicin were incubated at 28 °C until visible bacterial growth was observed. Subsequently, 50 µL of each culture was transferred into a fresh 5 mL 523 medium containing 1 mg/L rifampicin. This process was repeated with increasing rifampicin concentrations of 2, 5, 10, and finally 50 mg/L. Bacteria that grew in the highest rifampicin concentration (50 mg/L) were selected and stored for further use.

## Supporting information

Table S1. List of agrobacterial strains used in this study.

Table S2. List of agrobacterial concentrations used in the study in agroinfiltration experiments.

Table S3. List of primers used in the study.

Table S4. List of plasmids and sequences used in the study.

## Acknowledgments

We thank Dr. Kung-Ta Lee (Department of Biochemical Sciences, National Taiwan University, Taiwan), Dr. Jeff Chang (Department of Botany and Plant Pathology, Oregon State University, USA), and Dr. Shu-Yi Yang (Institute of Plant Biology, National Taiwan University, Taiwan) for sharing agrobacterial strains, Dr. Miin-Huey Lee (Department of Plant Pathology, National Chung Hsing University, Taiwan) for providing seeds of chili pepper. We also thank Shu-Ting Cho and Yu-Chen Lin (Institute of Plant and Microbial Biology, Academia Sinica, Taiwan) for technical assistance in genome sequencing and analysis, and the Genomic Technology Core (Institute of Plant and Microbial Biology, Academia Sinica, Taiwan) for Illumina sequencing library preparation. We thank Mark Youles (The Sainsbury Laboratory, UK) for sharing the plasmid pICH86922OD. This project was supported by the Academia Sinica Grand Challenge Program (AS-GC-111-L02 to EML, CHK, and CHW), the National Science and Technology Council of Taiwan (NSTC 112-2311-B-001-031 to CHK), and the intramural fund of the Institute of Plant and Microbial Biology, Academia Sinica.

## Author contributions

JCLA, FJG, EML, and CHW contributed to the conceptualization of the project. JCLA, FJG, ST, YSH, CYH, KTL contributed to data curation. JCLA, FJG, and ST contributed to the formal analysis. JCLA, FJG, and ST contributed to the investigation. YCW and YW contributed to the methodology. HXC contributed to resources. CHK, EML, and CHW supervised the research and acquired funding. CHW managed the project. JCLA, FJG, and ST contributed to data visualization. JCLA, and CHW wrote the initial manuscript draft. JCLA, CHK, EML, and CHW edited the manuscript.

## Data availability statement

The Illumina raw reads and the annotated genome sequence were deposited in NCBI GenBank and made publicly available under the accession number PRJNA721796 (https://www.ncbi.nlm.nih.gov/bioproject/PRJNA721796). Other data supporting the findings of this study are available within the article or as supporting information.

## Supporting Information

**Figure S1.** *R. rhizogenes* A4 transient expression efficiency is higher than the laboratory, biovar 1 strains AGL1 and EHA105.

**Figure S2.** *R. rhizogenes* A4 transient expression efficiency is higher than the biovar 2 strain A8196 and the rhizogenic strain ARqua1.

**Figure S3.** GFP fluorescence and protein accumulation are higher when transiently transforming *N. benthamiana* leaves using *R. rhizogenes* A4 compared to GV3101.

**Figure S4.** Undesirable traits were observed when using *R. rhizogenes* A4.

**Figure S5.** Confirmation of the removal of kanamycin resistance and introduction of rifampicin resistance in A4-derived strains.

**Figure S6.** Vacuum infiltration on 4-week-old *N. benthamiana* using the RUBY reporter results in high betalain production irrespective of the strain tested.

**Figure S7.** The putative type III secretion system of *R. rhizogenes* A4 does not contribute to *N. benthamiana* transient expression efficiency.

**Figure S8.** Sequences of csp22 peptides in GV3101 and A4 are highly conserved.

**Table S1.** List of agrobacterial strains used in this study.

**Table S2.** List of agrobacterial concentrations used in the study in agroinfiltration experiments.

**Table S3.** List of primers used in the study.

**Table S4.** List of plasmids and sequences used in the study.

