## Supplemental Figures S1-8 for "*Rhizobium rhizogenes* A4-derived strains mediate hyper-efficient transient gene expression in *Nicotiana benthamiana* and other solanaceous plants"

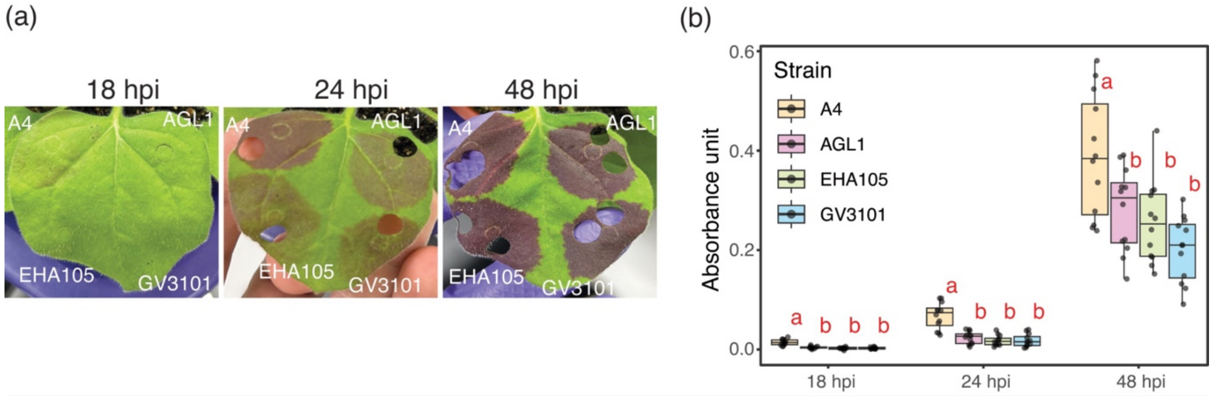


**Figure S1. *R. rhizogenes* A4 transient expression efficiency is higher than the laboratory, biovar 1 strains AGL1 and EHA105.** (a) Betalain accumulation on *N. benthamiana* leaves at 18, 24 and 48 hours post-infiltration comparing A4, AGL1, EHA105 and GV3101. (b) Betalain absorbance measurements were performed on the infiltration spots shown in (a) at three different timepoints (n=12). In box plots, the line inside the box indicates the median, the edges of the box represent the 25th and 75th percentiles, and the whiskers reach out to the most extreme data points, up to 1.5 times the interquartile range. Letters indicate statistically significant differences between treatments at each timepoint (Tukey's HSD test, $p < 0.05$).

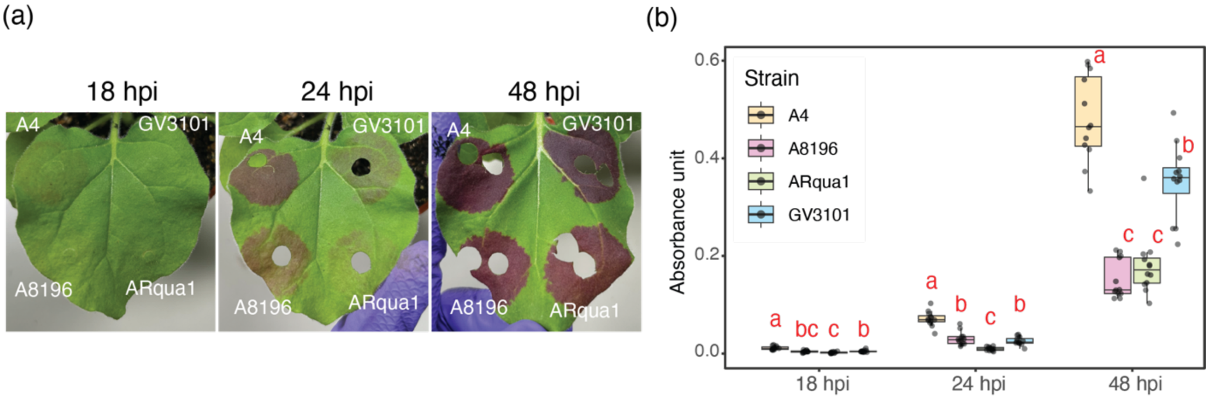


**Figure S2.** ***R. rhizogenes*** **A4 transient expression efficiency is higher than the biovar 2 strain A8196 and the rhizogenic strain ARqua1.** (a) Betalain accumulation on *N. benthamiana* leaves at 18, 24 and 48 hours post-infiltration comparing A4, GV3101, A8196 and ARqua1. (b) Betalain absorbance measurements were performed on the infiltration spots shown in (a) at three different timepoints (n = 12). In box plots, the line inside the box indicates the median, the edges of the box represent the 25th and 75th percentiles, and the whiskers reach out to the most extreme data points, up to 1.5 times the interquartile range. Letters indicate statistically significant differences between treatments at each timepoint (Tukey's HSD test, $p < 0.05$).

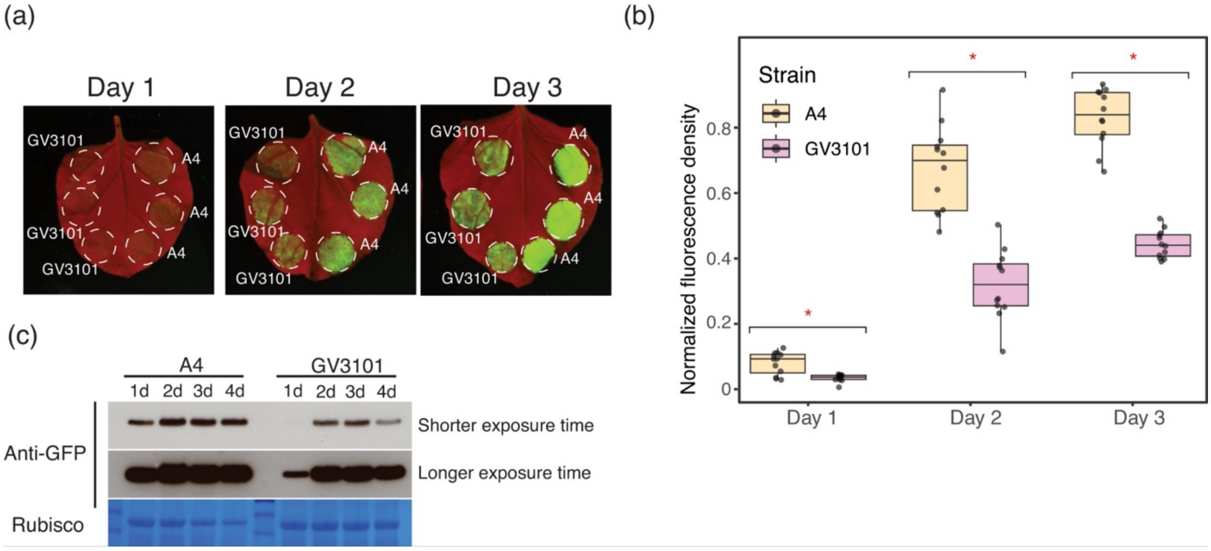


**Figure S3. GFP fluorescence and protein accumulation are higher when transiently transforming *N. benthamiana* leaves using A4 compared to GV3101.** (a) Imagining of GFP fluorescence using UVP ChemStudio PLUS on *N. benthamiana* leaves comparing A4 and GV3101 at three different timepoints. (b) GFP fluorescence quantification was performed on the infiltration spots shown in (a) at four different timepoints (n = 12). In box plots, the line inside the box indicates the median, the edges of the box represent the 25th and 75th percentiles, and the whiskers reach out to the most extreme data points, up to 1.5 times the interquartile range. * indicate statistically significant differences between treatments at each timepoint (Dunn's test, $p < 0.05$). (c) Western blot analysis of GFP driven by 35S promoter using A4 or G3101 strains. Samples were collected at 1, 2, 3 and 4 days (d) after infiltration. Protein in each infiltration spot was detected using an anti-GFP antibody. SimplyBlue SafeStain-staining of Rubisco was used as a loading control.

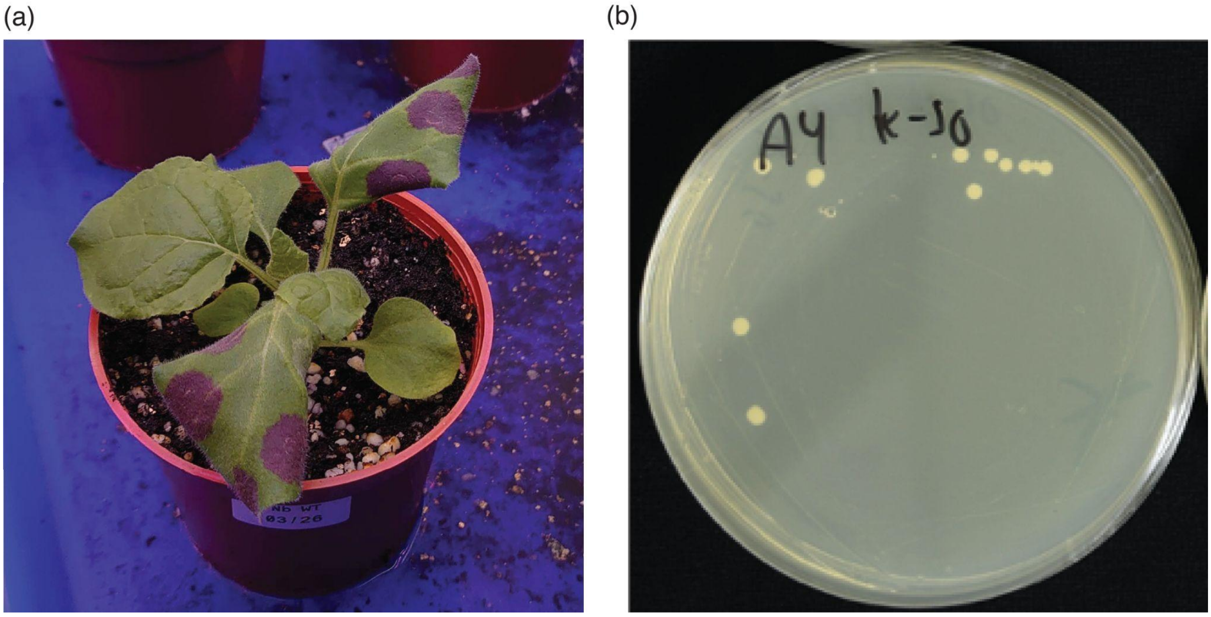


**Figure S4. Undesirable traits were observed when using *R. rhizogenes* A4.** (a) Agroinfiltration of *R. rhizogenes* A4 on *N. benthamiana* leaves leads to leaf curling. The photo was taken at 48 hours post-infiltration. (b) *R. rhizogenes* A4 develops spontaneous kanamycin-resistant colonies when grown on media containing 50 mg/L kanamycin. The photo was taken 3 days after plating and incubation at 28°C.

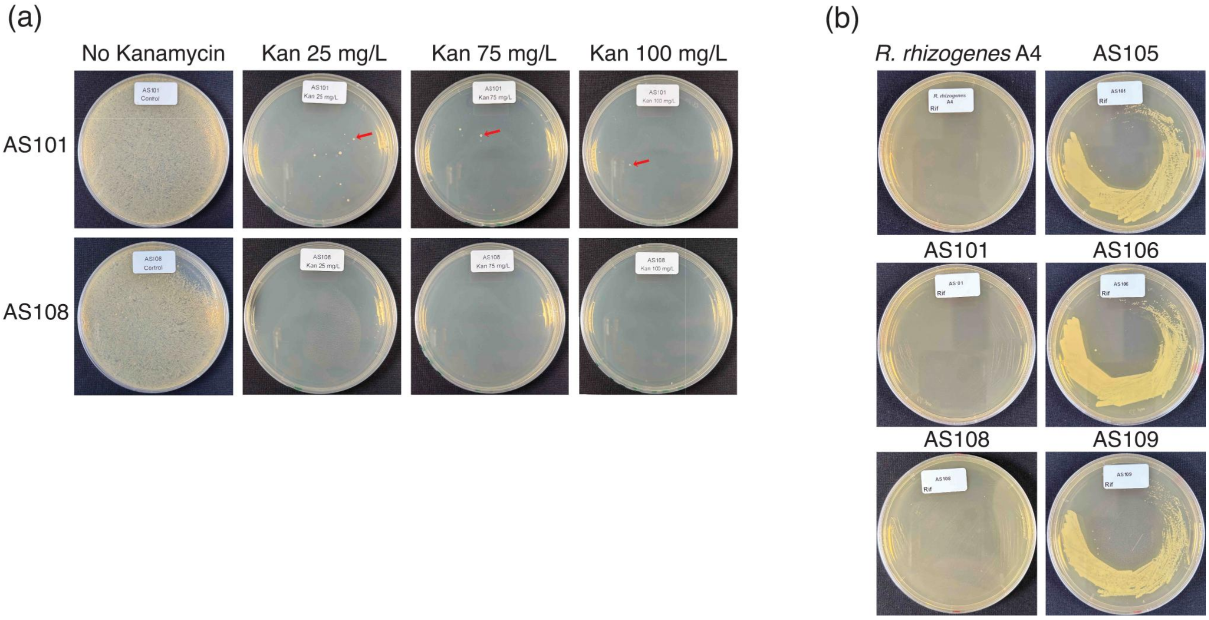


**Figure S5. Confirmation of the removal of kanamycin resistance and introduction of rifampicin resistance in A4-derived strains.** (a) Kanamycin sensitivity assay of the A4-derived strains AS101 and AS108. Bacteria were plated on 523 media containing 0, 25, 75 and 100 mg/L of kanamycin. Kanamycin-resistant colonies of AS101 appeared at any tested concentration (arrows), while AS108 (with a mutated kanamycin resistance gene) colonies did not appear. (b) Rifampicin resistance test comparing *R. rhizogenes* A4, AS101 and AS108 against their rifampicin-resistant derivatives AS105, AS106 and AS109. Bacteria was plated on media containing 50 mg/L rifampicin and photos were taken 3 days after plating and incubation at 28°C.

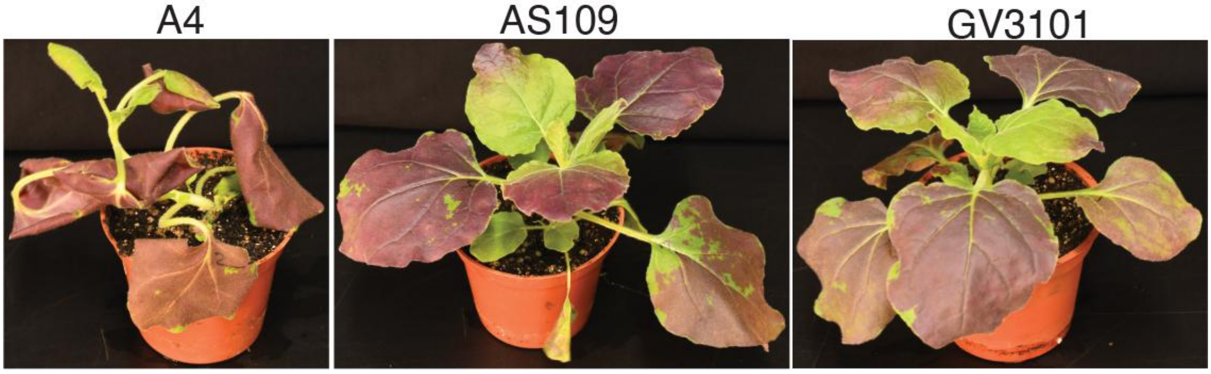


**Figure S6. Vacuum infiltration on 4-week-old *N. benthamiana* using the RUBY reporter results in high betalain production irrespective of the strain tested.** Transient expression efficiency on 4-week-old *N. benthamiana* leaves of A4, AS109 and GV3101 using vacuum infiltration and the RUBY reporter. Photos were taken at 3 days post-infiltration.

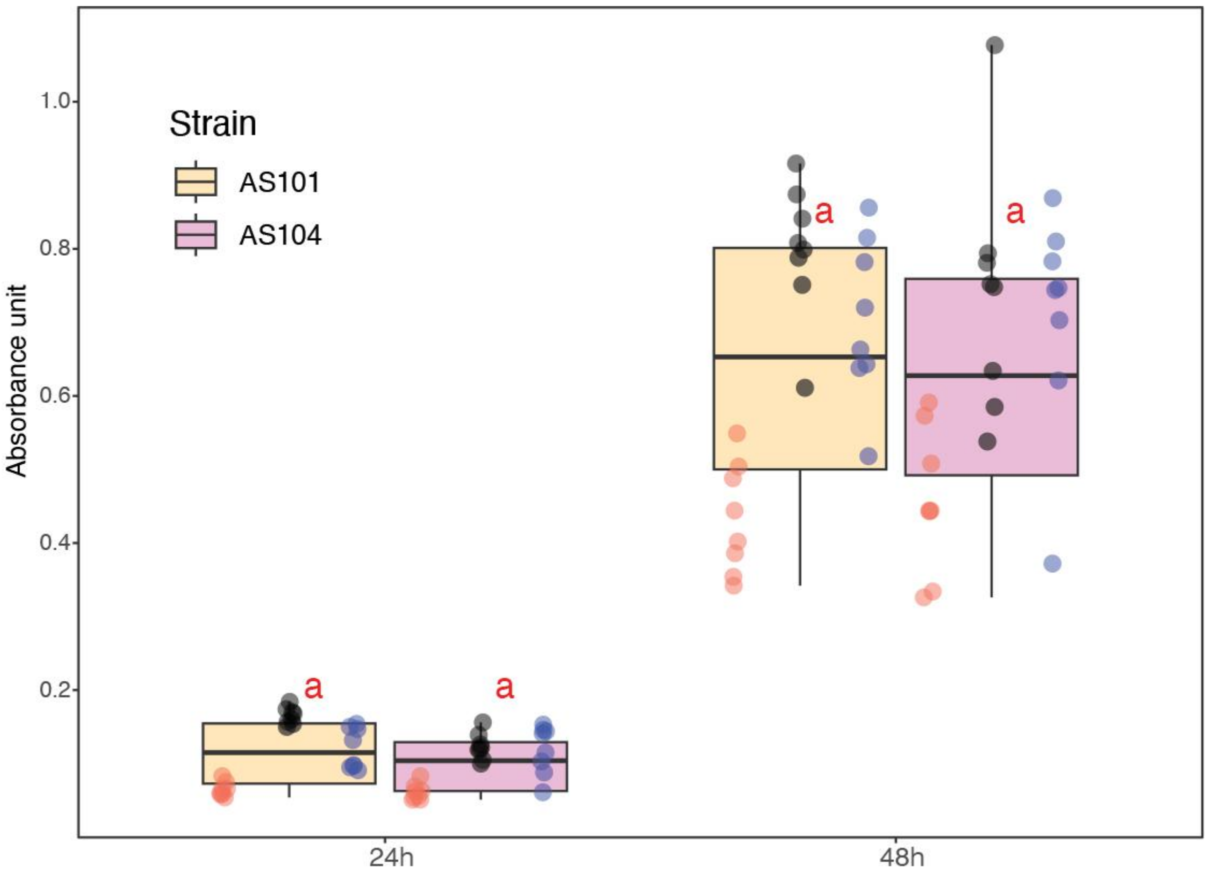


**Figure S7. The putative type III secretion system of *R. rhizogenes* A4 does not contribute to *N. benthamiana* transient expression efficiency.** Betalain absorbance was measured in infiltration spots on *N. benthamiana* leaves using AS101 (disarmed A4) or AS104 (A4 mutant with deleted putative T3SS gene cluster) at two timepoints. Each dot color represents an independent experiment ($n = 8$). In the box plots, the line within the box indicates the median, the box edges represent the 25th and 75th percentiles, and the whiskers extend to the most extreme data points within 1.5 times the interquartile range. Letters denote statistically significant differences between treatments at each timepoint (Tukey's HSD test, $p < 0.05$).

| Consensus (Wei et al., 2017) | AVGTVKWFNAEKGFGFITPDDG |
|---|---|
| GV3101\|EML485_19660 | ATGTVKFFAQDKGFGFITPDNG |
| A4\|EML492_09065 | TKGIVKFFNQDKGFGFITPDGG |
| A4\|EML492_27150 | PTGTVKFFNEDKGFGFITPENG |
| GV3101\|EML485_09180 | ETGTVKFFNTDKGFGFIKPDNG |
| A4\|EML492_10620 | ETGTVKFFNTDKGFGFIKPDKG |
| GV3101\|EML485_14370 | NTGTVKWFNATKGFGFIQPDNG |
| GV3101\|EML485_24560 | TTGTVKWFNSTKGFGFIQPDNG |
| A4\|EML492_25530 | TTGTVKWFNSTKGFGFIQPDDG |
| A4\|EML492_12325 | STGTVKWFNAAKGFGFIQPDDG |
| GV3101\|EML485_07265 | ITGVVKWFDVAKGFGFIVPDNG |
| A4\|EML492_07010 | ISGVVKWFDVAKGFGFIVPDNG |
| A4\|EML492_03075 | FNGIVKNFDLEKGYGFIQPTDG |
| GV3101\|EML485_12735 | ATGTVKWFNATKGYGFIQPDDG |

**Figure S8. Sequences of csp22 peptides in GV3101 and A4 are highly conserved.** Alignment of putative csp22 peptides identified in GV3101 and A4 according to the consensus sequence (Wei *et al.*, 2017).
